# Dynamics of alpha suppression index both modality specific and general attention processes

**DOI:** 10.1101/2022.11.16.516776

**Authors:** Grace M. Clements, Mate Gyurkovics, Kathy A. Low, Arthur F. Kramer, Diane M. Beck, M. Fabiani, G. Gratton

## Abstract

EEG alpha power varies under many circumstances requiring visual attention. However, mounting evidence indicates that alpha may not only serve visual processing, but also the processing of stimuli presented in other sensory modalities, including hearing. We previously showed that alpha dynamics during an auditory task vary as a function of competition from the visual modality (Clements et al., 2022) suggesting that alpha may be engaged in multimodal processing. Here we assessed the impact of allocating attention to the visual or auditory modality on alpha dynamics, during the preparatory period of a bimodal-cued conflict task. In this task, precues indicated the modality (vision, hearing) relevant to a subsequent reaction stimulus. This task afforded us the opportunity to assess alpha during modality-specific preparation and while switching between modalities. Alpha suppression following the precue occurred in all conditions indicating that it may reflect general preparatory mechanisms. However, we observed a switch effect when preparing to attend to the auditory modality, in which greater alpha suppression was elicited when switching to the auditory modality compared to repeating. No switch effect was evident when preparing to attend to visual information (although robust suppression did occur in both conditions). In addition, waning alpha suppression preceded error trials, irrespective of sensory modality. These findings indicate that alpha can be used to monitor the level of preparatory attention to process both visual and auditory information. These results support the emerging view that alpha band activity may index a general attention control mechanism used across modalities, at least vision and hearing.

## 1. Introduction

It is well recognized that EEG alpha power varies under many circumstances requiring visual attention in behaviorally beneficial ways (Gulbinaite et al., 2014; Heinrichs-Graham & Wilson, 2015; Thut et al., 2006). However, there is growing evidence that alpha may not only serve visual processing, but also the processing of stimuli presented in other sensory modalities, such as hearing (Banerjee et al., 2011; Elshafei et al., 2018; Fu et al., 2001). Indeed, we have shown that alpha dynamically engages during auditory tasks with and without visual input (Clements et al., 2022), suggesting that alpha may be involved in multimodal – or at least visual and auditory – processing and its associated cortical areas. It can therefore be hypothesized that a manipulation directing attention to the visual or auditory modality may also produce dynamic changes in alpha engagement. Currently, however, little is known about how alpha changes when attention is directed toward different modalities. Here we examined alpha dynamics during the preparatory period of a cued-conflict task, in which the precue informs participants about whether to respond on the basis of either the auditory or visual features of an upcoming bimodal reaction stimulus (thus engaging intermodal selective attention mechanisms).

Alpha band activity has been established as a mechanism by which selective attention within the visual domain is enabled. Previous work has observed alpha suppression after the presentation of visual stimuli (Yamagishi et al., 2005) and during the deployment of voluntary, cued visuospatial attention (Sauseng et al., 2005; Worden et al., 2000) indicating that alpha may be part of an active neural system supporting the allocation of attention to, and maintenance of, visual representations. Within such a framework, initial alpha suppression following a visual stimulus, either cue or target, likely helps select goal-relevant representations and focuses attention on the stimulus, in line with the theory proposed by Gratton (2018). This initial alpha suppression is primarily observed occipitally on the scalp and is thought to originate in early visual cortices (Romei et al., 2008; Yamagishi et al., 2005). Occipital alpha suppression has also been observed following auditory cues, to facilitate the deployment of visual selective attention (Fu et al., 2001). This work indicates that a modality other than vision can trigger changes in alpha, but it also still interprets alpha changes as related to *intramodal* (i.e., visual) selective attention mechanisms.

Another, more limited, line of research has also indicated that EEG oscillations in the alpha frequency might interact with auditory stimulus processing and be engaged in situations requiring auditory attention. Some investigators have successfully recorded alpha activity from (and localized generators in) primary auditory cortex (Lehtelä et al., 1997; Weisz et al., 2011 for review), but methodological challenges exist to noninvasively record “auditory alpha” due to the neuroanatomy of the primary auditory cortex.^1^ Given that the current study used scalp-recorded EEG measures, we focus on alpha that is produced in visual, or at most, multimodal regions.

Although cross-modal studies, in which multiple sensory streams are active, have demonstrated that alpha dynamics vary in early sensory cortices depending on whether the visual or auditory stream is selected (Elshafei et al., 2018; Keitel et al., 2013; Mazaheri et al., 2013; Saupe et al., 2009), these studies either used unimodal cues or measured alpha while continuous streams of unrelated auditory and visual stimuli were concurrently presented. Less has been done to elucidate alpha’s role in the *selection between modalities* during a preparatory interval in which *only* attention is manipulated. Whether and how alpha operates to allocate attention to a sensory stream and sustain that attentional focus over time when selection between two modalities is needed is still unknown.

Here we investigated the impact of a *bimodal*, informative, audiovisual precue on preparatory attention processes in which both sensory modalities receive the same information (i.e., participants see and hear the same informative precue). This allowed us to test whether scalp-recorded EEG alpha is a specific phenomenon, which operates only when instructed to attend to the visual modality, or whether instead, alpha suppression can be considered as a more general cognitive control phenomenon, which operates as a selection mechanism between two modalities (attend the visual stream vs. attend the auditory stream). If alpha is associated just with changes in visual attention processes, then alpha suppression should occur only when allocating attention to vision (but not to audition). If, instead, alpha is the manifestation of a general cognitive control mechanism, then alpha suppression should occur regardless of modality, but may vary in scalp topography depending on which modality attention is allocated to. Of course, a third possibility is that scalp-recorded EEG alpha may operate in both capacities to some extent, perhaps because it is produced both by unimodal visual areas and by multimodal regions. In this case, alpha suppression may be more pronounced when attention is directed to visual stimuli, but also be present to some extent when attention is directed to audition. Again, the scalp distribution of these two effects may be slightly different, reflecting the incomplete overlap in brain areas where the effect can be observed.

The bimodal cued-conflict paradigm allowed us to assess a second question: could alpha suppression be related to switching between attended modalities and is the effect of switching the same for the two modalities? Switching between attending to the visual and auditory modalities can be conceptualized as a general attention process in which attention to one modality is either maintained (in the case of modality repeat trials) or not (in the case of modality switch trials). As such, one may consider that changing attentional focus from one modality to another may involve greater alpha suppression – independent of the specific modalities involved – compared to a situation in which focus is maintained on the same modality across trials. In this case, one could argue that alpha suppression may be related to the operation of a general form of attention and cognitive control, rather than a more specific form related to certain task dimensions (e.g., sensory modality). It is also possible, that modality may interact with switching, where the difference in alpha suppression on switch versus repeat trials may vary as a function of modality. Indeed, there is evidence that both task-general and task-specific effects in neural dynamics exist and operate in conjunction with each other (Gratton et al., 2008; Mansfield et al., 2012).

Given its role in cognitive control, we also expected alpha to have consequences on participants’ performance, and we investigated whether or not this effect on behavior was dependent on modality. For this reason, we compared the alpha dynamics during the preparatory period for trials in which the participants responded correctly or incorrectly to the subsequent reaction stimulus. Should alpha suppression index general cognitive control operations, then any failure of that mechanism should impact performance in both attended modalities. If instead, alpha suppression indexes changes in visual attention, then failure of that operation should impact performance only when attending to visual information. Of course, an interaction effect is also possible, whereby failure to suppress alpha is associated with performance decrements in both modalities, but not to the same extent.

Thus, in the current study, we investigated whether alpha suppression (or lack thereof) is a general mechanism that reflects the occurrence of cognitive control operations, a more specific phenomenon that reflects changes in attentional focus within the visual modality, or both. In this context, three manipulations are considered: (a) modality that needs to be attended; (b) switching attentional focus between modalities; and (c) prediction of errors for either modality. The present data address all three questions and indicate that alpha may not merely be related to attentional changes in the processing of visual information, but a more general index of cognitive control observable across different modalities.

## 2. Method

### 2.1 Participants

A total of 59 young adults participated in this study. Nine participants were excluded due to equipment malfunction or experimenter error, resulting in a sample size of 50. Our criterion for inclusion was that participants retain 75% or more trials following EEG artifact detection. Two participants did not meet this criterion resulting in a final sample of 48 (*M*_*age*_ = 21.5, *SD* _*age*_ = 2.3, 91% female). All participants were native English speakers and reported themselves to be in good health with normal hearing and normal or corrected-to-normal vision, and free from medications that may directly affect the central nervous system. All were right-handed as assessed by the Edinburgh Handedness Inventory (Oldfield, 1971). All participants signed informed consent, and all procedures were approved by the University of Illinois at Urbana-Champaign Institutional Review Board.

### 2.2 Task and Procedures

Each trial began with a white fixation cross presented in the middle of a black screen for 1600 ms. This was followed by a bimodal precue presentation for 400 ms, consisting of either the letter A (for auditory) or the letter V (for visual). The same letter was presented in *both* the auditory (70 dB SPL) and visual modalities, and served as a precue, instructing the participant about which modality to attend on the upcoming reaction stimulus. After a 1600 ms fixation interval, the reaction stimulus, consisting of one auditory and one visual letter, was presented for 400 ms. These could be either the letter “I” or the letter “O” (**Figure 1**). On half the trials the *same* letter was presented in both modalities (*congruent condition*: audiovisual “I” or audiovisual “O”), whereas on the other half a different letter was presented in each modality (*incongruent condition*: one I and one O, differing by modality). Thus, participants could respond above chance to the reaction stimulus only if they had processed the precue information. The to-be-attended modality indicated by the precue was randomized (.5 probability) across trials. This allowed us to sort trials based on whether the relevant modality matched that of the previous trial. We labeled the trials in which the relevant modality remained the same as in the previous trial the “repeat” condition; those in which it changed, the “switch” condition. Participants were instructed to respond to the letter in the cued modality as rapidly as possible, while still maintaining high accuracy. Right- and left-hand button presses were mapped to the two letters, and the stimulus-response mapping was counterbalanced across participants. Participants were instructed to attend to the fixation cross, bimodal precue and subsequent bimodal reaction stimulus regardless of the cued modality. Participants were specifically instructed not to close their eyes or look away on auditory trials and were monitored via closed-circuit video to ensure compliance. We collected 20 experimental blocks, with 24 trials per block, for a total of 480 trials. The overall design included 16 conditions: two switch levels (switch, repeat); two attended modalities (auditory, visual); two levels of congruency (congruent, incongruent); two test letters (I, O) mapped to a right/left button press. Prior to the experimental blocks, study participants were also given 8 blocks of practice (16 trials per block), with the first two blocks presented at half-speed (i.e., longer precue-to-target and target-to-precue intervals) in order to introduce the task, which was typically perceived as difficult.

**Figure 1:**
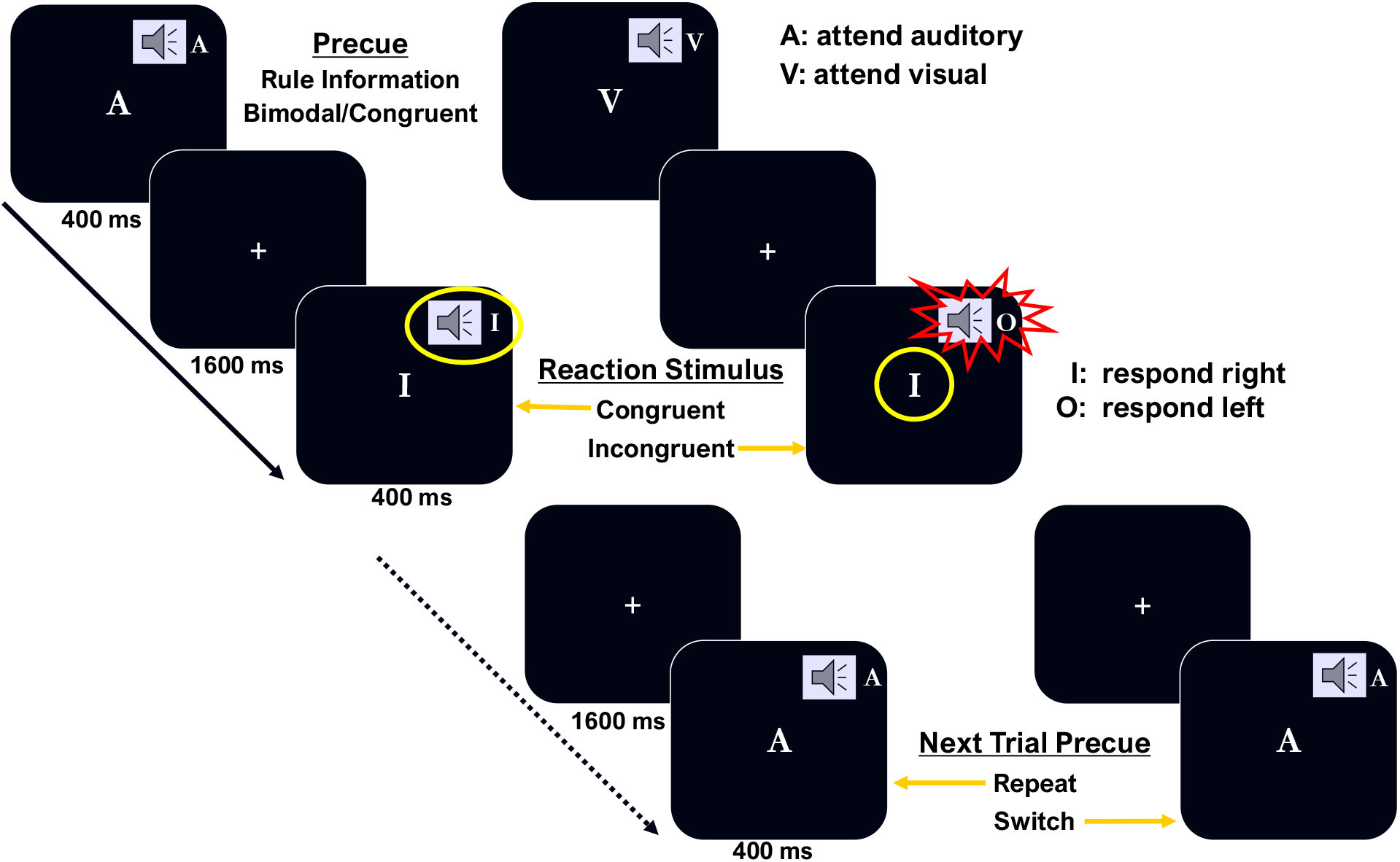
Trial Schematic. A bimodal precue was presented for 400 ms, followed by a delay of 1600 ms. Then a bimodal reaction stimulus was presented for 400 ms, and could be either congruent or incongruent. The next trial began with a precue 1600 ms after the offset of the reaction stimulus.

This experiment was part of a larger study involving the effect of video game training on cognitive function. Only data from session one of the larger study (i.e., collected prior to video game training) is included in the current analyses.

### 2.3 EEG Recording and Preprocessing

Electroencephalographic (EEG) activity was recorded with electrode caps fitted with 23 tin electrodes (Electro-Cap International, Inc) in the standard 10-20 electrode configuration (Jasper, 1958). A Grass Model 12 amplifier with a bandpass setting of .01 to 30 Hz was used for data recording with a sampling rate of 100 Hz. Scalp electrodes were referenced to an electrode placed over the left mastoid and re-referenced offline to the average of the two mastoids. Eye movements and blinks were monitored with bipolar recordings from the left and right outer canthi of the eyes and above and below the left eye. Offline processing of EEG was performed using the EEGLAB Toolbox (version: 2021.1, Delorme & Makeig, 2004) and custom MATLAB 2021a scripts (The MathWorks Inc., Natick, MA, USA). The data were epoched into 3500 ms segments relative to precue onset, including 1500 ms of EEG recording before and 2000 ms after precue onset. Epochs with amplifier saturation were discarded (less than .01% of all trials). Ocular artifacts were corrected using the procedure described in Gratton et al. (1983), based on the bipolar EOG recordings. After eye movement correction, epochs with voltage fluctuations exceeding 200 *μ*V were excluded from further analysis to minimize the influence of any remaining artifactual activity. As a reminder, our criterion for inclusion was that participants had to have 75% or more of their trials retained following artifact detection. If more than 25% of a participant’s epochs were marked for rejection, they were visually inspected to determine if one or two faulty electrodes were the cause. If so, their traces were replaced with the interpolated traces of the neighboring electrodes and reprocessed to regain the lost epochs.

Time-frequency representations of the data were then derived using Morlet wavelet convolution with MATLAB scripts modified from Cohen (2014) and according to the recommendations in Keil et al. (2022). Epoched data were fast Fourier transformed (FFT) and multiplied by the fast Fourier transform of Morlet wavelets of different frequencies. Morlet wavelets are complex sine waves tapered by a Gaussian curve. Thirty logarithmically spaced wavelets between 3 and 30 Hz were used. The width of the Gaussian taper ranged between 3-10 cycles and logarithmically increased as a function of frequency in order to balance the tradeoff between temporal and frequency precision.

An inverse Fourier transform was applied to the product of the FFT’d wavelets and the FFT’d data, and power values were computed by calculating the modulus of the complex values from the iFFT (i.e., by squaring the length of this complex vector at each time point). To reduce edge artifacts during convolution, each epoch was tripled in length by using reflections on either side of the original epoch, such that the original epoch was sandwiched between two reflected versions of itself. Following time-frequency derivation, the reflected epochs were removed restoring the original length of 3500 ms.

Power values were baseline corrected using condition-specific subtractive baselining. A condition-specific baseline was used because the content of the previous trial had meaning (switch vs. repeat) which could result in lingering unique activity in the baseline period. We have previously shown that, compared to divisive baselining, subtractive baselining minimizes the potential of Type I errors that might occur because of the aperiodic, 1/*f* component of power spectra in certain situations (Clements et al., 2021, 2022; Gyurkovics et al., 2021). The power in the baseline period (−750 to −250 ms) was thus subtracted from power values across the whole epoch, frequency by frequency. A baseline period of this length provides adequate temporal resolution in the alpha frequency band (8-12 Hz), our primary interest. It also minimizes the influence of edge effects and reduces any impact of activity from the previous trial contaminating our estimate of baseline activity and as a result, the baseline correction procedure. The final 250 ms of each epoch was also ignored for analysis, to reduce the possibility of activity generated by the reaction stimulus being temporally smeared into the pre-stimulus time window during wavelet convolution. Thus, the analyzed epoch length was 2500 ms: from 750 ms before the precue to 1750 ms after.

### 2.4 Conditions of Interest

The EEG data were sorted in two ways for analysis purposes. First, to investigate the effects of cued modality and task-switching on alpha activity in the preparatory period, time-frequency power on correct trials was collapsed across congruency and response hand. This was done because the congruency and response hand for the upcoming reaction stimulus are not known during the preparatory interval and collapsing increases statistical power to test the main manipulations of interest: modality of attention and task-switching. Only correct trials were used in this analysis because participants did not make enough errors to have adequate numbers of trials in the modality and switch bins. We did not combine correct responses and errors because we predicted that there would be differences between correct and incorrect preparation (see Introduction). Sorting trials this way resulted in a 2 (Modality) × 2 (Switching) repeated measures design. Second, all error trials (collapsed across all conditions) were compared with a subset of correct trials selected randomly within-subjects, such that each participant had the same number of correct and error trials. Participants made relatively few errors, so collapsing this way was necessary. This pairing allowed us to test whether alpha power fundamentally differed during adequate preparation, leading to correct responses, and inadequate preparation, resulting in errors.

### 2.5 Statistical Approach

Given our primary aim of investigating changes in preparatory alpha activity, we chose to analyze the time-frequency data at a set of posteriorly located electrodes (Pz, Oz, P3, P4, O1, O2) that closely match those used in Clements et al. (2022). These electrodes were further subdivided into a parietal (Pz, P3, P4) and occipital (Oz, O2, O2) subset to investigate potential differences in alpha activity at different scalp locations during preparation. To conduct a targeted analysis of alpha, we chose to only analyze the alpha power time series from 8-12 Hz. This frequency band matches with the alpha effects reported in Clements et al. (2022). The *difference* between alpha power time series’ from two conditions was compared to zero, indicating no difference between conditions, within nine consecutive 200 ms time-windows (0-1750 ms) at parietal and occipital electrode subsets. First, the interaction timeseries between modality and switch on alpha power was assessed with this method by comparing the modality effect on switch trials (Visual Switch – Auditory Switch) to the modality effect on repeat trials (Visual Repeat – Auditory Repeat) using the following subtraction:

Interaction = (Visual Switch – Auditory Switch) – (Visual Repeat – Auditory Repeat) Significant interactions were followed up by tests of simple main effects of switch and modality. Preparatory alpha power on correct and error trials was similarly assessed (with the subtraction: Correct - Error).

Because of wavelet convolution, the data were temporally smoothed. As such each millisecond of data does not reflect a unique measurement of power. At 10 Hz, we estimated the temporal integration window used in wavelet convolution to be approximately 200 ms, making it an ideal time window to assess alpha oscillatory changes across the preparatory period. Bootstrapped confidence intervals using 1000 bootstrap samples were calculated for the alpha difference time series in each of the 9 time intervals. A 99% confidence interval, Bonferroni corrected for multiple comparisons (100*(1 – 0.05/9) = 99.4%) was calculated for each time point within each of the 9 time windows. Note that this is a conservative correction given that time windows are not independent. Only time windows for which the 99% bootstrapped CIs did not contain zero for the entire 200 ms time period were considered to show significant differences between conditions. The dashed lines along, and perpendicular to, the alpha time series in the upcoming figures denote the average activity in each 200 ms time period. To help visualize alpha activity patterns, time-frequency maps and scalp topographies were also generated and are presented.

Accuracy and reaction time (RT) were used to assess behavioral performance during this task. Responses made with the incorrect hand and timeouts were not included in the accuracy data and false alarms were removed from the RT data prior to analysis. To match the time-frequency analysis, data were collapsed over response hand, target congruency, and test letters (I, O) resulting in a 2 (Modality) × 2 (Switching) repeated measures ANOVA, which was conducted in R (version 4.0.2; R Core Team, 2020). Normality was checked using the Shapiro-Wilk test and confirmed by examining Q-Q plots.

## 3. Results

### 3.1 Behavior

The mean reaction time (RT) and accuracy data can be found in **Table 1**. Participants were more accurate on attend-visual compared to attend-auditory trials, *F*(1, 47) = 14.105, *p* < .001. There was also a main effect of switch, such that participants were more accurate on repeat trials compared to switch trials, *F*(1, 47) = 17.254, *p* < .001. The interaction was not significant, *F*(1, 47) = 0.096, *p* = .758. The RT data paralleled the accuracy data with a main effect of modality, in which attend-visual trials were responded to more quickly than attend-auditory trials, *F*(1, 47) = 17.183, *p* < .001. Repeat trials were also responded to more quickly than switch trials, *F*(1, 47) = 8.312, *p* < .01. The interaction was not significant, *F*(1, 47) = 0.156, *p* = .695. The modality main effects could be due to a more automatic sight-to-hand than listening-to-hand response mapping, as previously reported (Gladwin & De Jong, 2005; Stephan & Koch, 2010, 2011, 2016). Compatible mappings between stimulus and response (either visual-manual or auditory-vocal) tend to prime the selection of the response in the compatible modality. This task does not require a vocal response to an auditory stimulus and so would not benefit from the compatible mapping, which might explain the slower and decreased performance on the attend-auditory trials relative to the attend-visual trials.

**Table 1:**
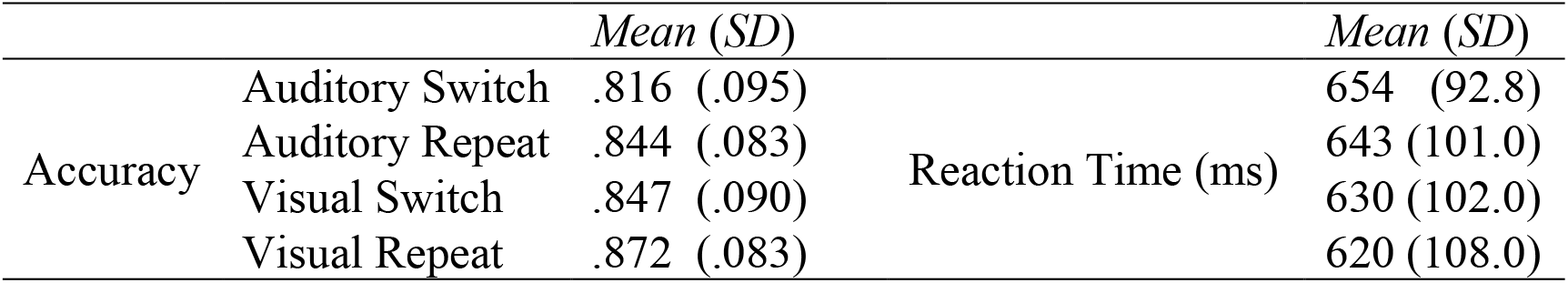
Behavioral Data - Descriptive Statistics.

### 3.2 Alpha Suppression

We first examined the main effects of modality and switching. Visual trials produced more alpha suppression than auditory trials, which was reliable from 600-1000 ms at parietal sites and from 600-1200 and 1400-1600 ms at occipital sites. For the main effect of switch, the only reliable difference was between 600-800 ms at parietal electrodes (see **Supp. Figures 1 & 2**). These main effects, however, were largely superseded by interactions of switch and modality during these same time windows, as well as during the 200 ms immediately following the precue.

The early interaction (0-200 ms) occurred at parietal electrodes and was driven by alpha *enhancement* (see **Figure 2** insets) on visual switch trials. Follow up analyses for each modality separately indicate a significant switch effect for visual (**Figure 3)**, but not for auditory (**Figure 4)** during this early period. A possible interpretation of the initial alpha enhancement is contamination from evoked potentials elicited by the precue, which might differ in terms of scalp distribution when attention is deployed to the visual or auditory modality. Therefore, these findings should be interpreted with some caution.

**Figure 2:**
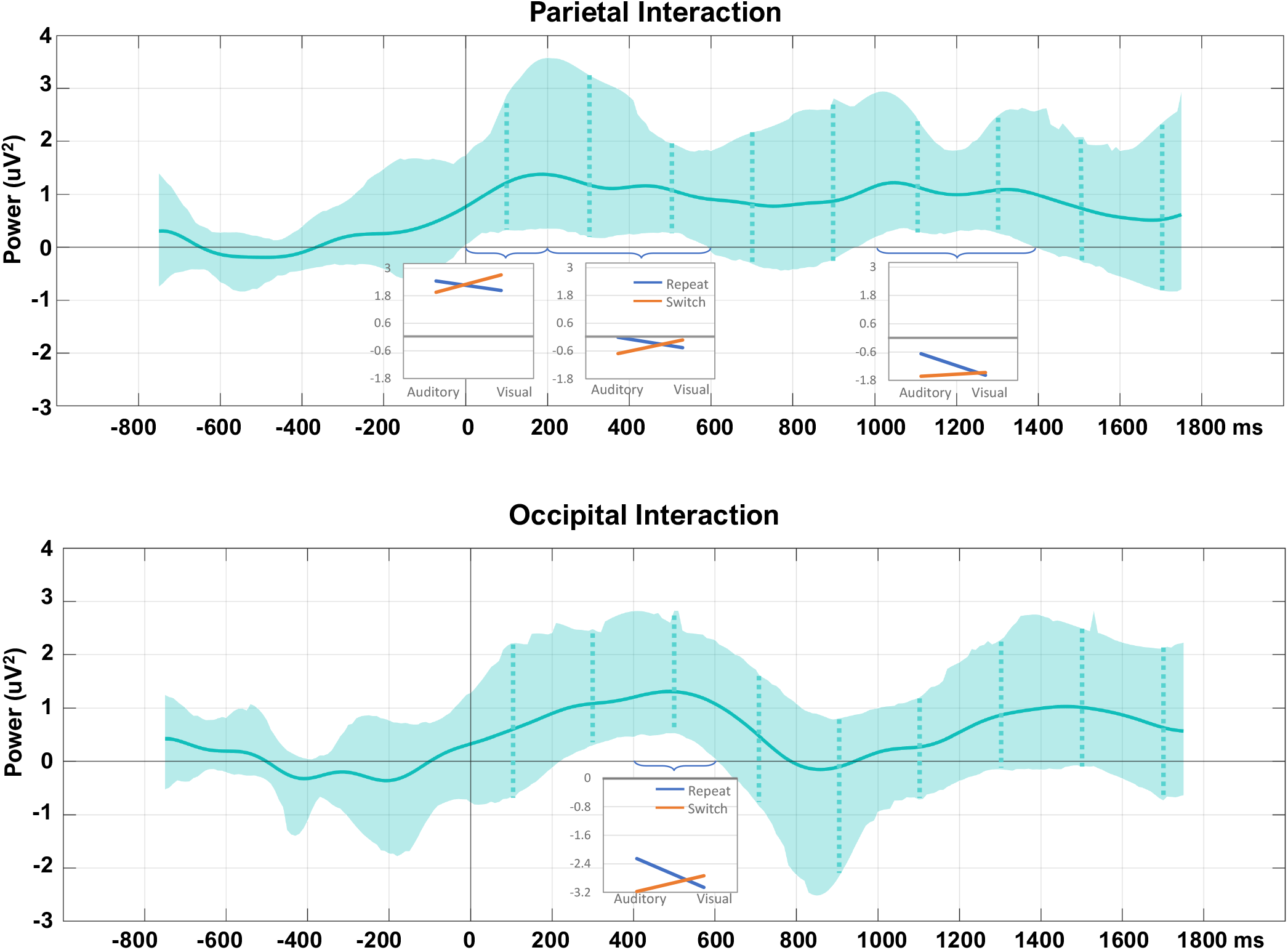
Interaction between switch and modality on alpha activity (8-12 Hz), calculated as the modality difference on repeat trials (Visual Repeat – Auditory Repeat) subtracted from the modality difference on switch trials (Visual Switch – Auditory Switch) at both parietal (top) and occipital (bottom) electrodes. Insets illustrate the interaction at each of the significant time windows and the inset scales are the same within a subplot. Shading indicates 99% bootstrapped confidence intervals. The dotted vertical lines indicate the center of each 200 ms analytic interval.

**Figure 3:**
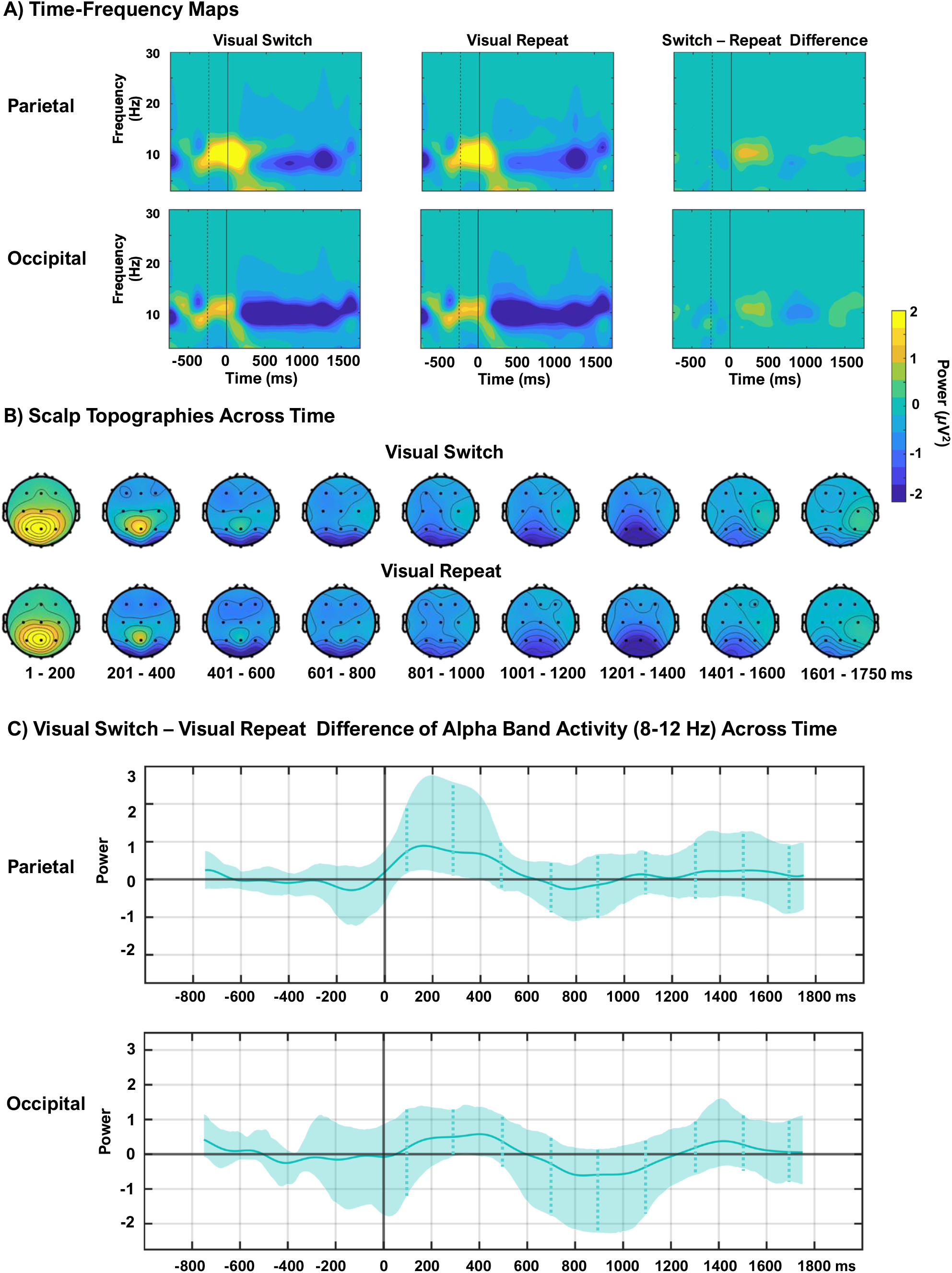
Comparison of visual trials. Time-frequency maps (A) of the preparatory period for attend-visual switch responses, attend-visual repeat responses, and switch minus repeat response differences. The dotted vertical line indicates the end of the baseline period, the solid vertical line indicates precue onset. Note: statistical testing of the difference map was not performed, and this panel is displayed for visualization only. Statistics were limited to the alpha time series, in line with hypotheses. Scalp topographies (B) across the preparatory period for attend-visual switch (top) and attend-visual repeat trials (bottom). A and B are on the same color scale. Difference waveforms (C) of the alpha timeseries (8-12 Hz) with 99% bootstrapped confidence intervals indicate no significant differences. The dotted vertical lines indicate the center of each 200 ms analytic interval. In A and C, the top row includes activity from the parietal electrodes, the bottom includes activity from the occipital electrodes

**Figure 4:**
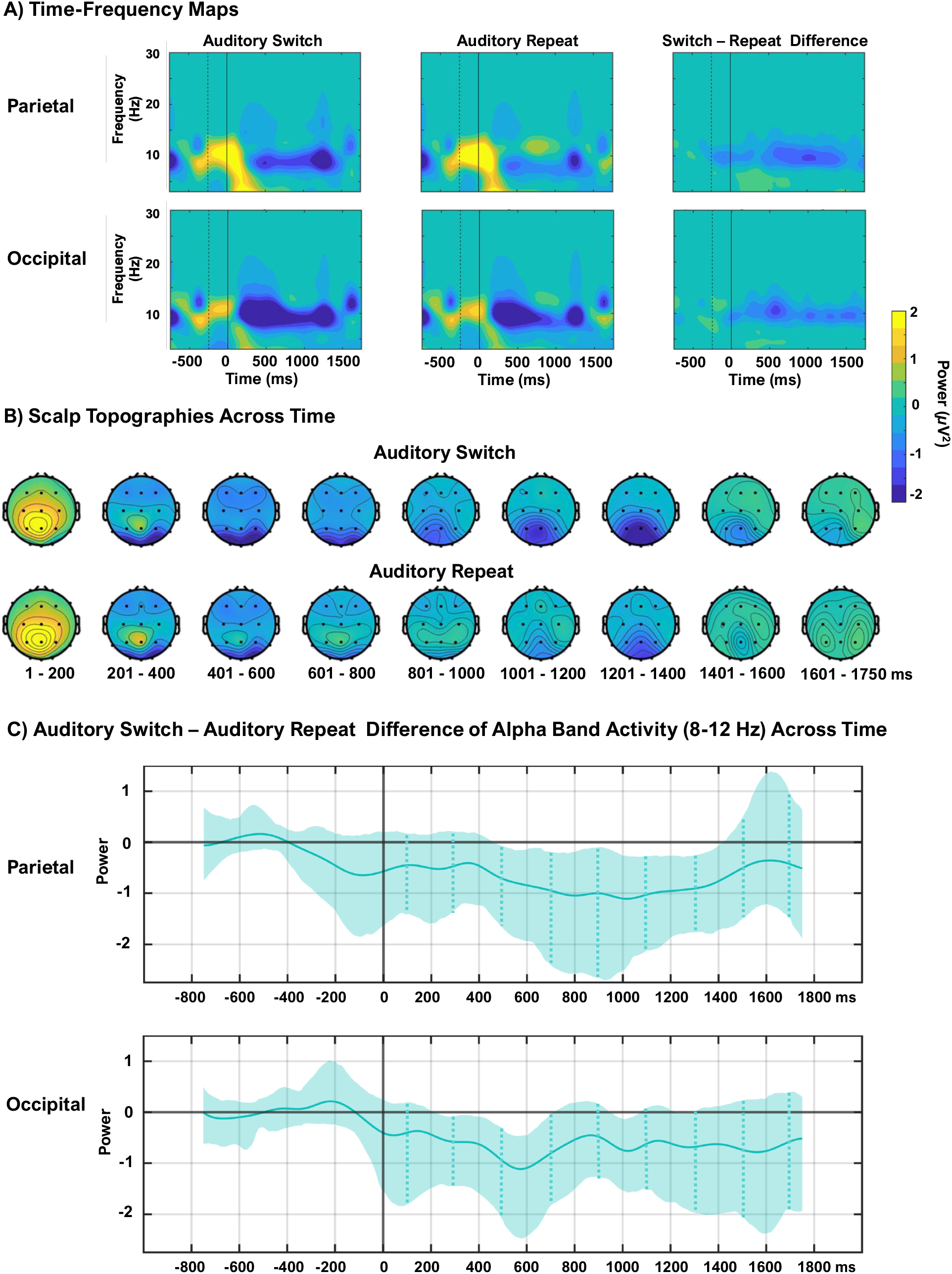
Comparison of auditory trials. Time-frequency maps (A) of the preparatory period for attend-auditory switch responses, attend-auditory repeat responses, and the switch minus repeat response differences. The dotted vertical line indicates the end of the baseline period, the solid vertical line indicates precue onset. Note: statistical testing of the difference map was not performed, and this panel is displayed for visualization only. Statistics were limited to the alpha time series, in line with hypotheses. Scalp topographies (B) across the preparatory period for attend-auditory switch (top) and attend-auditory repeat trials (bottom). A and B are on the same color scale. Difference waveforms (C) of the alpha timeseries (8-12 Hz) with 99% bootstrapped confidence intervals indicate that at parietal electrodes, a sustained significant difference exists from 600 - 1400 ms after the precue. The dotted vertical lines indicate the center of each 200 ms analytic interval In A and C, the top row includes activity from the parietal electrodes, the bottom includes activity from the occipital electrodes.

In contrast, the interactions in later intervals, which were reliable at parietal sites, 200-600 and 1000-1400 ms, and at occipital sites at 400-600 ms, all reflect differences in alpha *suppression*. Follow up analyses of switch effects separately for each modality indicate that, on visual trials, there were no differences in alpha suppression for switch versus repeat trials (**Figure 3**), whereas auditory trials showed greater alpha suppression for switch compared to repeat (**Figure 4**). This finding is consistent with the interpretation that alpha reflects more general cognitive control operations, as it is present on both attend auditory and attend visual trials. However, the lack of switch effects on visual trials and the overall main effect of greater alpha suppression on visual compared to auditory trials indicate a more robust coupling (or tuning) between visual attention demands and alpha suppression; that is, visual attention is associated with robust alpha suppression regardless of whether the trial requires a repetition of attend visual or a switch to visual from auditory. The complementary simple effects of modality, separately for repeat and switch trials can be seen in **Supp. Figure 3 & 4**.

### 3.3 Effect of Accuracy

Lastly, we analyzed whether alpha differed in the preparatory period preceding errors compared to correct responses, indicative of inadequate preparation. As a reminder, the number of correct trials was restricted to match the number of error trials at the individual subject level. As can be seen in the time-frequency maps in **Figure 5A**, correct trials had more sustained alpha suppression than errors. However, the difference in alpha suppression between correct and error trials was most evident and significantly less than zero only at parietal electrodes beginning at 1000 ms and continuing until the end of the measurement period (**Figure 5C**), statistically confirming the larger suppression visualized in the time-frequency maps and scalp topographies prior to correct trials. Compared to correct trials, the precue interval preceding errors was characterized by a less pronounced alpha suppression, followed by a reestablishment and enhancement of alpha at parietal electrodes, as can be seen by positive power values (in yellow) in the pre-subtraction time-frequency maps (**Figure 5A**) and scalp topographies (**Figure 5B**).

**Figure 5:**
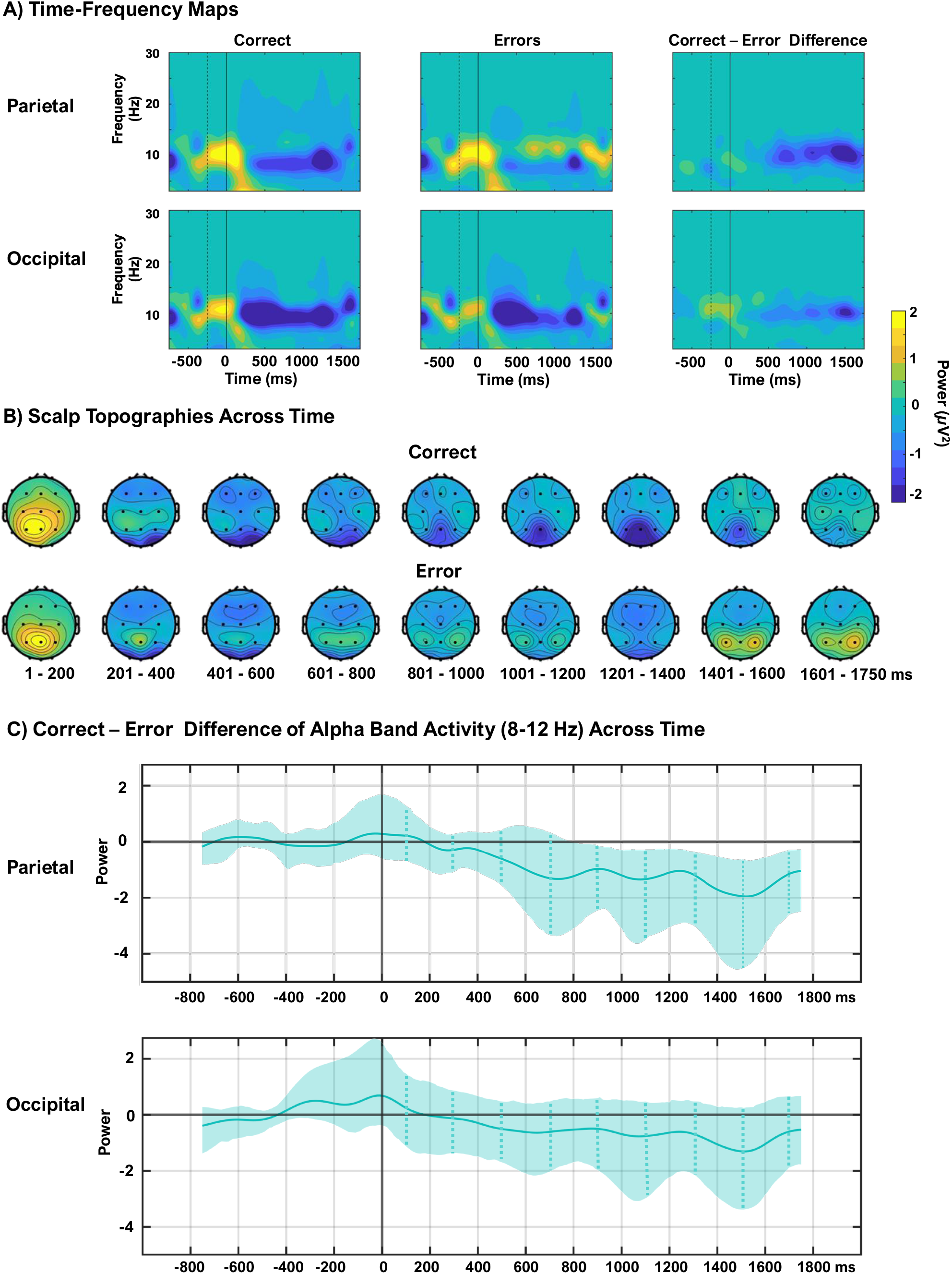
Comparison of correct and error trials. Time-frequency maps **(A)** of the preparatory period for correct responses, error responses, and for correct minus error response differences. The dotted vertical line indicates the end of the baseline period, the solid vertical line indicates precue onset. Note: statistical testing of the difference map was not performed, and this panel is displayed for visualization only. Statistics were limited to the alpha time series, in line with hypotheses. Scalp topographies **(B)** across the preparatory period for correct (top) and error trials (bottom). A and B are on the same color scale. Difference waveforms **(C)** of the alpha time series (8-12 Hz) with 99% bootstrapped confidence intervals indicate that at parietal electrodes, a significant difference begins at 800 ms after the precue. The dotted vertical lines indicate the center of each 200 ms analytic interval. In A and C, the top row includes activity from the parietal electrodes, the bottom includes activity from the occipital electrodes.

The waning parietal suppression and reestablishment of alpha late in the epoch was predictive of errors, suggesting that parietally-maximal alpha suppression must be sustained to fully process the upcoming reaction stimulus, and that the reemergence of alpha late in the preparatory period interrupts successful target processing.

We additionally compared attend-auditory to attend-visual errors and did not find evidence of a modality difference on error trials. Parietal alpha activity preceding errors in both modalities looked remarkably similar (time-frequency plots, **Supp. Figure 5A**; scalp topographies **Supp. Figure 5B**) and the bootstrapped alpha time series difference was not significant (**Supp. Figure 5C**), indicating that errors resulted from a mis-engaged alpha mechanism, regardless of modality. The occipital responses preceding attend-visual and attend-auditory errors were also remarkably similar and not statistically different (**Supp. Figure 5**). These data suggest that all errors, regardless of attended modality or switch-condition, were associated with similar alpha dynamics, and provide further evidence that alpha suppression helps maintain task-representations via general cognitive control processes.

### 3.4 Other Considerations

We assessed whether the parietal and occipital alpha suppression conditional differences were distinguishable from each other by directly comparing the conditional difference waves for the two scalp locations using the bootstrap procedure described above. Effectively, this assesses whether there is an interaction between scalp location and conditional difference (attend-visual vs. attend-auditory; switch vs. repeat; correct vs. error). These direct comparisons did not reveal any interactions between scalp location and condition for alpha suppression. Chi-square tests of independence also failed to reveal any reliable interactions between scalp location and condition.

To rule out the possibility that participants who performed poorly on the task were driving the alpha effects reported above, we performed a median split of participants based on their performance (accuracy) on incongruent trials to generate two groups. Then, we analyzed the “high-performers” and “low-performers” time-frequency responses on correct trials only (collapsed over response hand.) As expected, the low-performers (*M* = 0.78, *SD* = 0.06) performed significantly worse on the task than high-performers (*M* = 0.90, *SD* = 0.04), *t*(48) = −8.035, *p* < .001. To correctly respond on incongruent trials, participants must have processed the precue and directed their attention to the cued modality to resolve interference on the reaction stimulus. High- and low-performers had very similar patterns of conditional differences, indicating that the effects reported above did not result from a different alpha preparatory pattern in one group versus the other (**Supp. Figure 6** for modality effects; **Supp. Figure 7** for switch effects). However, it should be noted that these tests compared one half of the subjects to the other (each group *n* = 24) resulting in reduced statistical power to assess these effects.

## 4. Discussion

Our findings indicate that alpha suppression may be a mechanism indexing general cognitive control operations above and beyond attentional changes in the processing of visual information. Some level of alpha suppression occurred following the audiovisual precue in all conditions, but this level deepened when attending the visual modality and with increasing task-difficulty. Alpha suppression in response to attend-visual precues was not only deeper than attend-auditory precues but it was more robust to modality switching (i.e., it was equivalent in both switch and repeat trials). This pattern is consistent with a visual bias in alpha suppression during preparation; that is, the system may be predisposed to attend to the visual modality. This bias is consistent with faster RTs and higher accuracy in the visual modality. However, alpha suppression also occurred on attend-auditory trials, and was faster and more pronounced when auditory precues occurred after a visual trial than after an auditory trial. In fact, it emerged more quickly than when the precue indicated switching to the visual modality (which showed more extended early alpha enhancement), suggesting an important role for alpha suppression in auditory attention as well as visual. Additionally, a lack of alpha suppression predicted poorer performance, indicating that these alpha dynamics played a functional role in attention processing. Together, these effects suggest that the presence of alpha suppression indicates that attention has been effectively deployed and shows that it engages variably on auditory trials, with the most suppression occurring when switching away from vision.

Generally, the presence of alpha signaled that attention had been deployed in preparation for an upcoming reaction stimulus in either modality. However, we detected an interaction between modality and switch, illustrating a more complex picture in which directing attention to the auditory modality resulted in more flexible engagement of alpha compared to attending to the visual modality. If the magnitude of alpha suppression reflects an index of cognitive control operations, then switching to auditory attention requires more control (at least within the first 400 ms) than any form of visual attention. It may be that participants are predisposed to maintain high levels of visual selective attention control because with open eyes, visual attention *could* be directed anywhere at any time. This would result in robust alpha suppression in all visual attention situations, as reported here. High levels of visual selective attention, and concurrently high levels of alpha suppression, may be the default processing mode and obligatory.

Although the system may be set to process visual information, precues directing attention to the auditory modality engendered suppression. This indicates that alpha suppression at posterior electrodes occurs not only when attention is directed toward particular locations of the visual field (e.g., Thut et al., 2006; Yamagishi et al., 2005), but also when attention is directed away from the visual and toward the auditory modality. This provides evidence for a general role of alpha suppression that serves multiple modalities.

The interaction between modality and switch occurred at an early and a late interval and was particularly evident at parietal locations. The early interval (0-600 ms) had two components: an alpha enhancement from 0-200 ms, characterized by larger alpha on visual switch vs. auditory switch trials, and an alpha suppression from 200-600 ms, characterized by larger suppression on auditory switch vs. visual switch trials. The initial enhancement may reflect the time-frequency instantiation of the N1-P2 complex (which itself is in the alpha range) or prestimulus activity temporally smeared due to wavelet convolution, and therefore it is difficult to interpret. The subsequent suppression is dominated by rapid suppression on auditory-switch trials (most evident on scalp topographies for Visual and Auditory Switch, **Supp. Figure 4**) indicating that switching to auditory attention engages more quickly than switching to visual attention. At least two explanations could be posited for this. First, rapid alpha suppression in response to switching-to the auditory modality may be required to override the modality-bias of the visual attention system as reported previously (Colavita, 1974; Lukas, Phillip, & Koch, 2009; Posner, Nissen, & Klein, 1976). This condition may require more attention control and so control operations are enabled quickly to account for this. Second, because the auditory system is anatomically more compact than the visual system, sensory information reaches primary auditory cortex faster than it reaches primary visual cortex (Chatrian et al., 1960; Creel, 2019; Picton et al., 1974). In turn, this may result in more rapid engagement of auditory attention processes than visual ones. Finally, the later interaction effect (1000-1400 ms) followed the pattern seen in the earlier 200-400 ms time interval (see **Figure 2** insets), consistent with the interpretation that sustained alpha suppression serves a general role of cognitive control for multiple modalities.

We did not find evidence for scalp topography differences. It may be that effects between scalp locations are small and with the current sample size we are unable to detect them. We may be able to detect differences between parietal and occipital alpha using source localization methods (Xie & Richards, 2022) or fast optical imaging, as Parisi et al. (2020) have done during a visuospatial attention paradigm. It may also be that alpha suppression is such a large signal that it dominates all recordings from posterior locations. Additionally, with our data we cannot rule out the fact that a similar selective attention mechanism may be occurring in lateral, temporal cortices that have a bias toward the auditory modality (as Frey et al., 2014 and Weisz et al., 2020 have shown) which we cannot detect with scalp-recorded EEG.

Lastly, a sustained parietal alpha suppression reflected a process that was predictive of behavior. Greater parietal alpha suppression occurred on correct trials compared to error trials, regardless of attended modality or switch condition, and once it began it was sustained until the end of the epoch (800-1750 ms). If the sustained suppression waned and alpha rebounded or increased, participants were more likely to commit an error on the upcoming reaction stimulus than if the suppression persisted, suggesting that this suppression was required to maintain the information contained within the precue in higher-order attention control regions, also indicating a general role of alpha suppression to enable cognitive control operations. Sustained parietal suppression was needed to do the task correctly, maintain a general attentional focus (no differences existed between attend-auditory and attend-visual error trials), and successfully execute a response. Some limitations should be pointed out. This study only investigated the impact of auditory and visual attention on alpha dynamics and therefore our conclusions only relate to the integration of those two modalities. There are, of course, three other senses in which expectation and preparation may occur in the brain. Somatosensory preparatory processes are easiest to test, which would bolster our claims about the role of alpha suppression as a general index of cognitive control processes across modalities. Given that this study was conducted exclusively with open eyes, we did not observe post-stimulus alpha enhancement, which has been previously reported with eyes closed (Clements et al., 2022). Some elegant experiments could be designed to assess unimodal auditory attention with closed eyes – in which alpha enhancement would be expected – or bimodal auditory and somatosensory attention with the eyes open or closed. These would show the impact of engaging or disengaging the visual system on preparatory processing in other modalities. It is also worth noting that these effects could reflect, in part, variability in non-oscillatory, broadband (1/*f*) activity. The steepness of 1/*f* activity has been reported to vary after presentation of attentionally relevant auditory stimuli (Gyurkovics et al., 2022). However, an in-depth investigation of broadband activity is beyond the scope of this paper.

In conclusion, the current study provides evidence that alpha suppression may be an index of general cognitive control operations, useful beyond the visual modality. Alpha suppression following a bimodal informative precue occurred in all conditions and generally indicated that preparatory attention had been deployed. However, we observed a switch effect when preparing to attend to auditory information, which was not evident when preparing to attend to visual information (although robust suppression did occur in both conditions). This, along with waning alpha suppression preceding error trials, which was not sensitive to modality, indicate that alpha can be used to monitor level of attention/preparation not only for processing visual information but also for processing auditory information. These results support the emerging view that alpha band activity may index a general attention control mechanism used across modalities, at least vision and hearing.

## Acknowledgments

This work was supported by NIA grant RF1AG062666 to G. Gratton and M. Fabiani and the Office of Naval Research Multidisciplinary University Research Initiative (MURI) grant N00014-07-1-1913 to Arthur F. Kramer. We acknowledge Jason Agran for help with data collection, and Andreas Keil, Sepideh Sadaghiani, and Walter Boot for helpful comments. An earlier version of this manuscript was included in the dissertation of the first author in partial fulfillment of the requirement of the Ph.D. degree at the University of Illinois at Urbana-Champaign.

## Footnotes

1 Non-invasively, MEG must be used to detect auditory alpha because Heschl’s gyrus in primary auditory cortex is deeply folded. The activity generated there is undetectable with EEG because the non-parallel electrical dipoles cancel each other out. Primary auditory cortex is also much smaller than primary visual cortex, rendering occipitally generated alpha more prominent in M/EEG recordings (Weisz, 2011). Electrocorticography has been used to invasively measure alpha activity generated in primary auditory cortex (e.g., Nourski et al., 2021).

## Notes

### Competing Interest Statement

The authors have declared no competing interest.

