## Supplemental Figures for "Dynamics of alpha suppression index both modality specific and general attention processes"

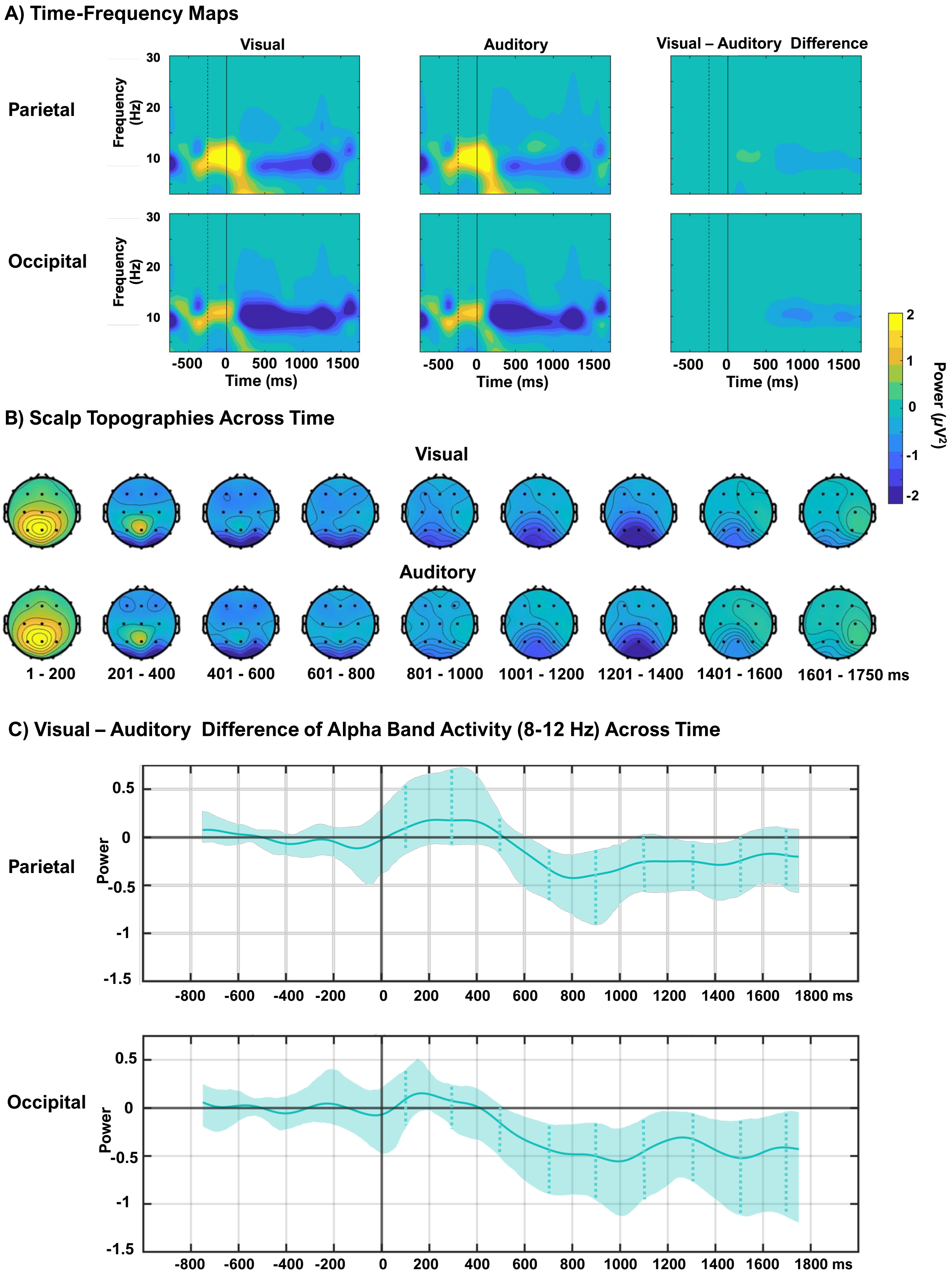

**Supp. Figure 1**: **Main Effect of Modality**. Time-frequency maps (A) of the preparatory period for attend-visual responses, attend-auditory responses, and the visual minus auditory response differences. The dotted vertical line indicates the end of the baseline period, the solid vertical line indicates precue onset. Note: statistical testing of the difference map was not performed, and this panel is displayed for visualization only. Statistics were limited to the alpha time series, in line with hypotheses. Scalp topographies (B) across the preparatory period for attend-visual (top) and attend-auditory trials (bottom). A and B are on the same color scale. Difference waveforms (C) of the alpha timeseries (8-12 Hz) with 99% bootstrapped confidence intervals indicate that at occipital electrodes, a sustained significant difference begins at 600 ms after the precue. The parietal effect is less sustained. The dotted vertical lines indicate the center of each 200 ms analytic interval. In A and C, the top row includes activity from the parietal electrodes, the bottom includes activity from the occipital electrodes.

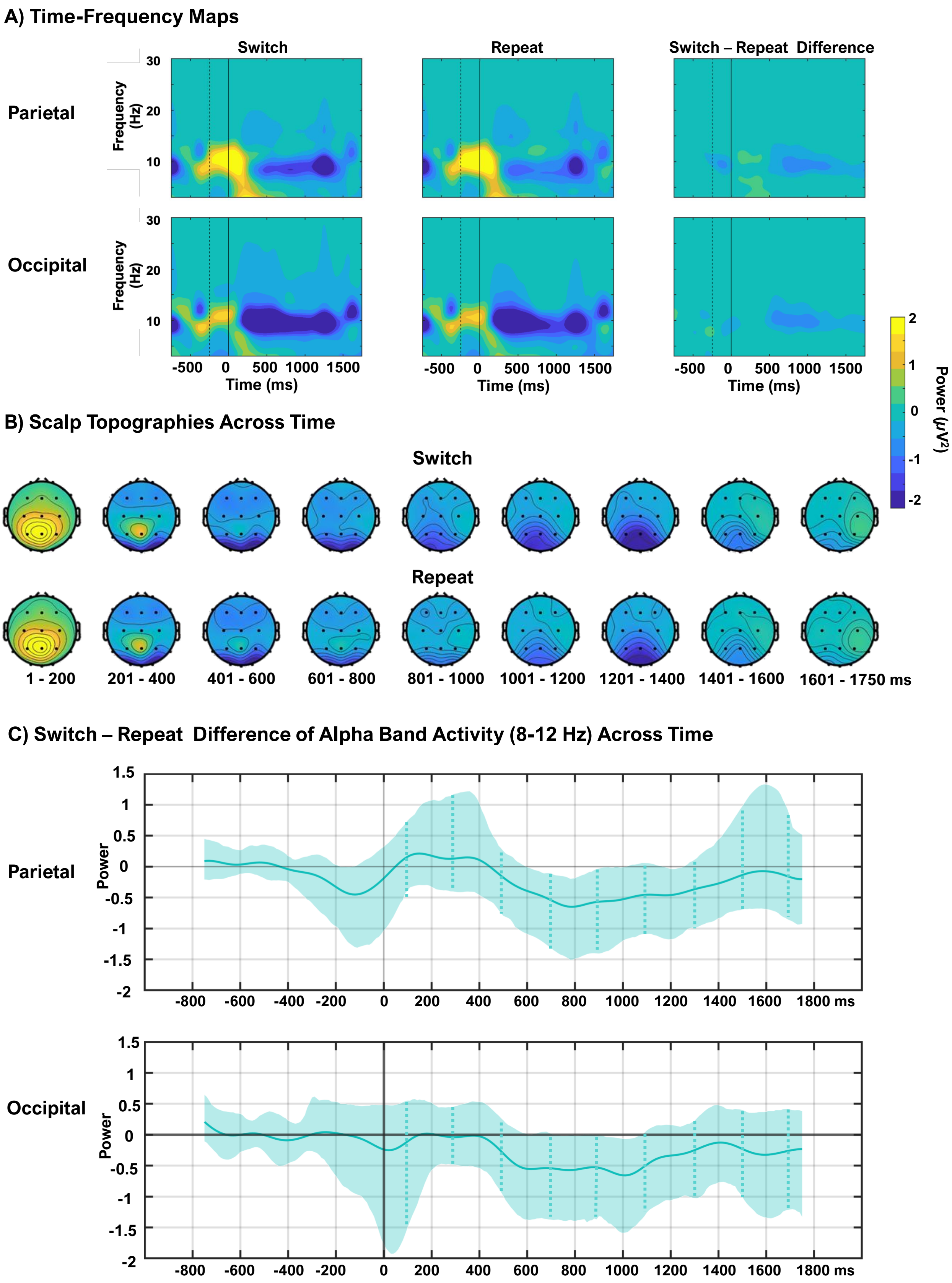
A) Time-Frequency Maps
Switch
Repeat
Switch – Repeat Difference
Parietal
Occipital
Frequency (Hz)
30
20
10
-500 0 500 1000 1500
Time (ms)
Power (μV²)
2
1
0
-1
-2
B) Scalp Topographies Across Time
Switch
Repeat
1 - 200
201 - 400
401 - 600
601 - 800
801 - 1000
1001 - 1200
1201 - 1400
1401 - 1600
1601 - 1750 ms
C) Switch – Repeat Difference of Alpha Band Activity (8-12 Hz) Across Time
Parietal
Occipital
Power
1.5
1
0.5
0
-0.5
-1
-1.5
-2
-800 -600 -400 -200 0 200 400 600 800 1000 1200 1400 1600 1800 ms

**Supp. Figure 2**: **Main Effect of Switch**. Time-frequency maps (A) of the preparatory period for switch responses, repeat responses, and the switch minus repeat differences. The dotted vertical line indicates the end of the baseline period, the solid vertical line indicates precue onset. Note: statistical testing of the difference map was not performed, and this panel is displayed for visualization only. Statistics were limited to the alpha time series, in line with hypotheses. Scalp topographies (B) across the preparatory period for attend-visual (top) and attend-auditory trials (bottom). A and B are on the same color scale. Difference waveforms (C) of the alpha timeseries (8-12 Hz) with 99% bootstrapped confidence intervals indicate a transient significant difference from 600-800 ms at parietal electrodes, which was not evident at occipital locations. The dotted vertical lines indicate the center of each 200 ms analytic interval. In A and C, the top row includes activity from the parietal electrodes, the bottom includes activity from the occipital electrodes.

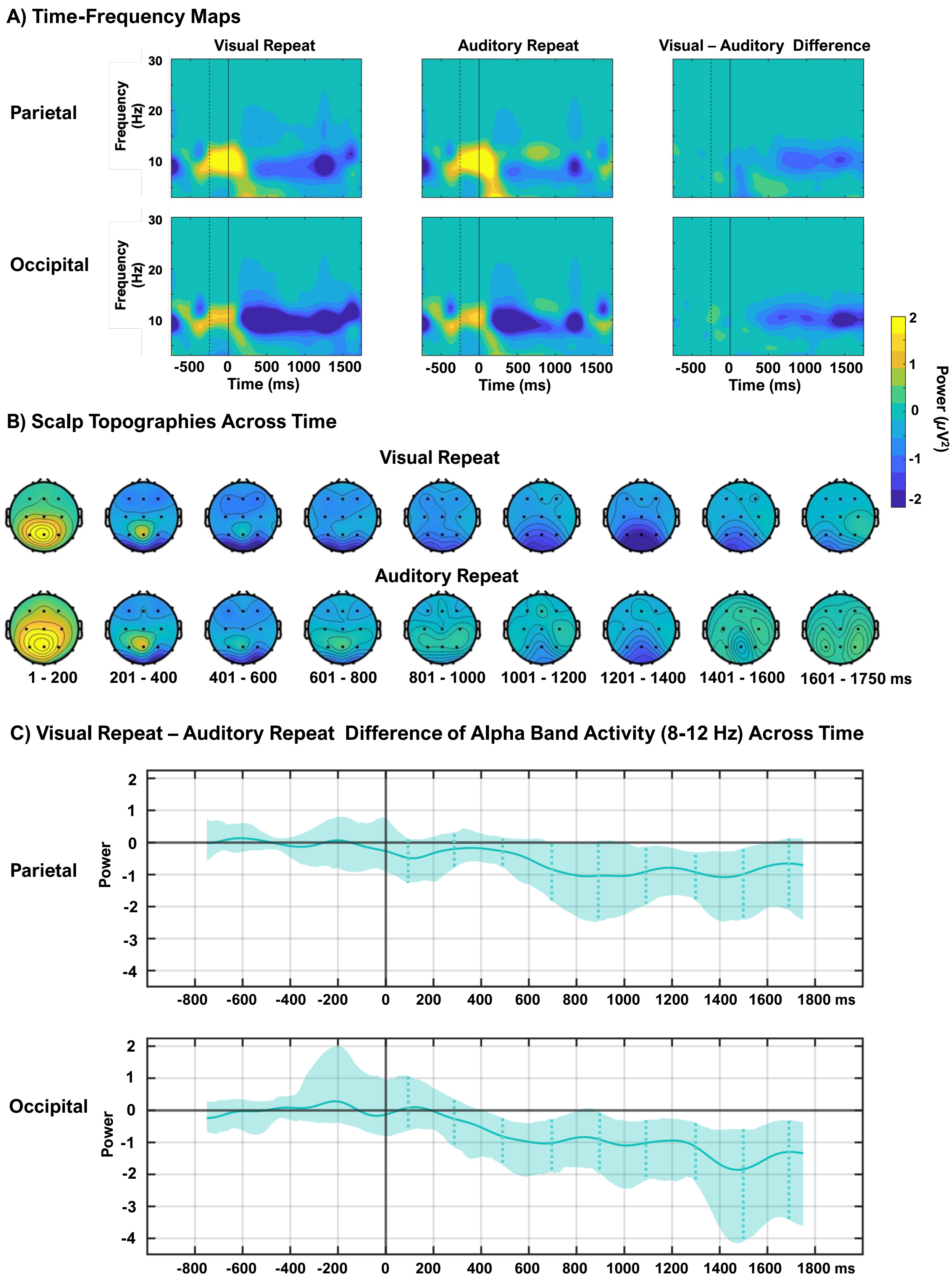
A) Time-Frequency Maps
Visual Repeat
Auditory Repeat
Visual – Auditory Difference
Parietal
Occipital
Frequency (Hz)
30
20
10
-500 0 500 1000 1500
Time (ms)
Power (μV²)
2
1
0
-1
-2
B) Scalp Topographies Across Time
Visual Repeat
Auditory Repeat
1 - 200
201 - 400
401 - 600
601 - 800
801 - 1000
1001 - 1200
1201 - 1400
1401 - 1600
1601 - 1750 ms
C) Visual Repeat – Auditory Repeat Difference of Alpha Band Activity (8-12 Hz) Across Time
Parietal
Occipital
Power
2
1
0
-1
-2
-3
-4
-800 -600 -400 -200 0 200 400 600 800 1000 1200 1400 1600 1800 ms

**Supp. Figure 3**: **Comparison of repeat trials**. Time-frequency maps (A) of the preparatory period for attend-visual repeat responses, attend-auditory repeat responses, and the visual repeat minus auditory repeat response differences. The dotted vertical line indicates the end of the baseline period, the solid vertical line indicates precue onset. Note: statistical testing of the difference map was not performed, and this panel is displayed for visualization only. Statistics were limited to the alpha time series, in line with hypotheses. Scalp topographies (B) across the preparatory period for attend-visual repeat (top) and attend-auditory repeat trials (bottom). A and B are on the same color scale. Difference waveforms (C) of the alpha timeseries (8-12 Hz) with 99% bootstrapped confidence intervals indicate that at occipital electrodes, a sustained significant difference begins at 400 ms after the precue. The parietal effect is less sustained. The dotted vertical lines indicate the center of each 200 ms analytic interval. In A and C, the top row includes activity from the parietal electrodes, the bottom includes activity from the occipital electrodes.

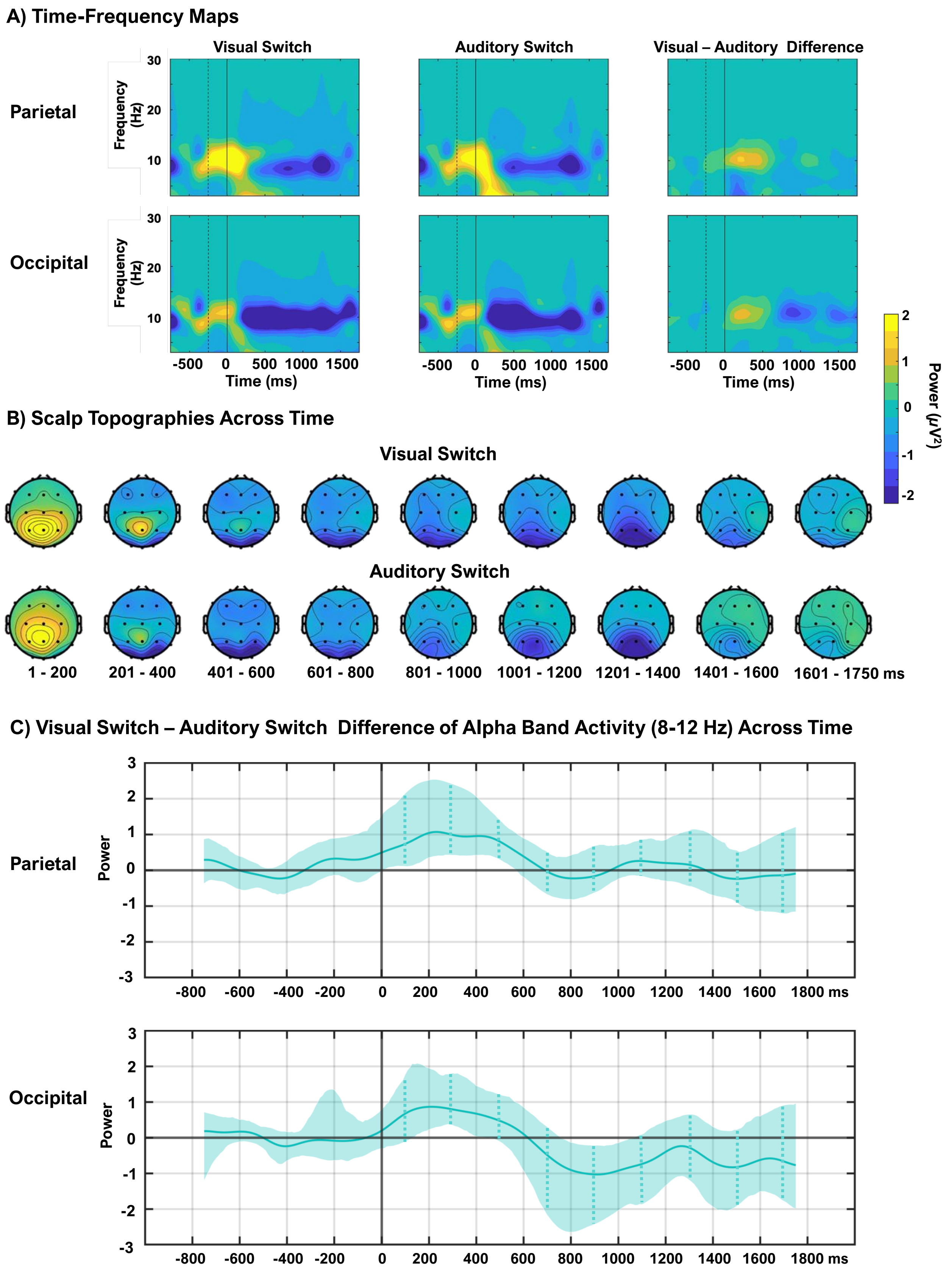

A) Time-Frequency Maps
Visual Switch
Auditory Switch
Visual – Auditory Difference
Parietal
Occipital
Frequency (Hz)
30
20
10
-500
0
500
1000
1500
Time (ms)
Power (μV²)
2
1
0
-1
-2
B) Scalp Topographies Across Time
Visual Switch
Auditory Switch
1 - 200
201 - 400
401 - 600
601 - 800
801 - 1000
1001 - 1200
1201 - 1400
1401 - 1600
1601 - 1750 ms
C) Visual Switch – Auditory Switch Difference of Alpha Band Activity (8-12 Hz) Across Time
Parietal
Occipital
Power
3
2
1
0
-1
-2
-3
-800
-600
-400
-200
0
200
400
600
800
1000
1200
1400
1600
1800 ms

**Supp. Figure 4**: **Comparison of switch trials**. Time-frequency maps (A) of the preparatory period for attend-visual switch responses, attend-auditory switch responses, and the visual switch minus auditory switch response differences. The dotted vertical line indicates the end of the baseline period, the solid vertical line indicates precue onset. Note: statistical testing of the difference map was not performed, and this panel is displayed for visualization only. Statistics were limited to the alpha time series, in line with hypotheses. Scalp topographies (B) across the preparatory period for attend-visual switch (top) and attend-auditory switch trials (bottom). A and B are on the same color scale. Difference waveforms (C) of the alpha timeseries (8-12 Hz) with 99% bootstrapped confidence intervals indicate that at parietal and occipital electrodes, a transient significant difference exists from 200-400 ms after the precue. The dotted vertical lines indicate the center of each 200 ms analytic interval. In A and C, the top row includes activity from the parietal electrodes, the bottom includes activity from the occipital electrodes.

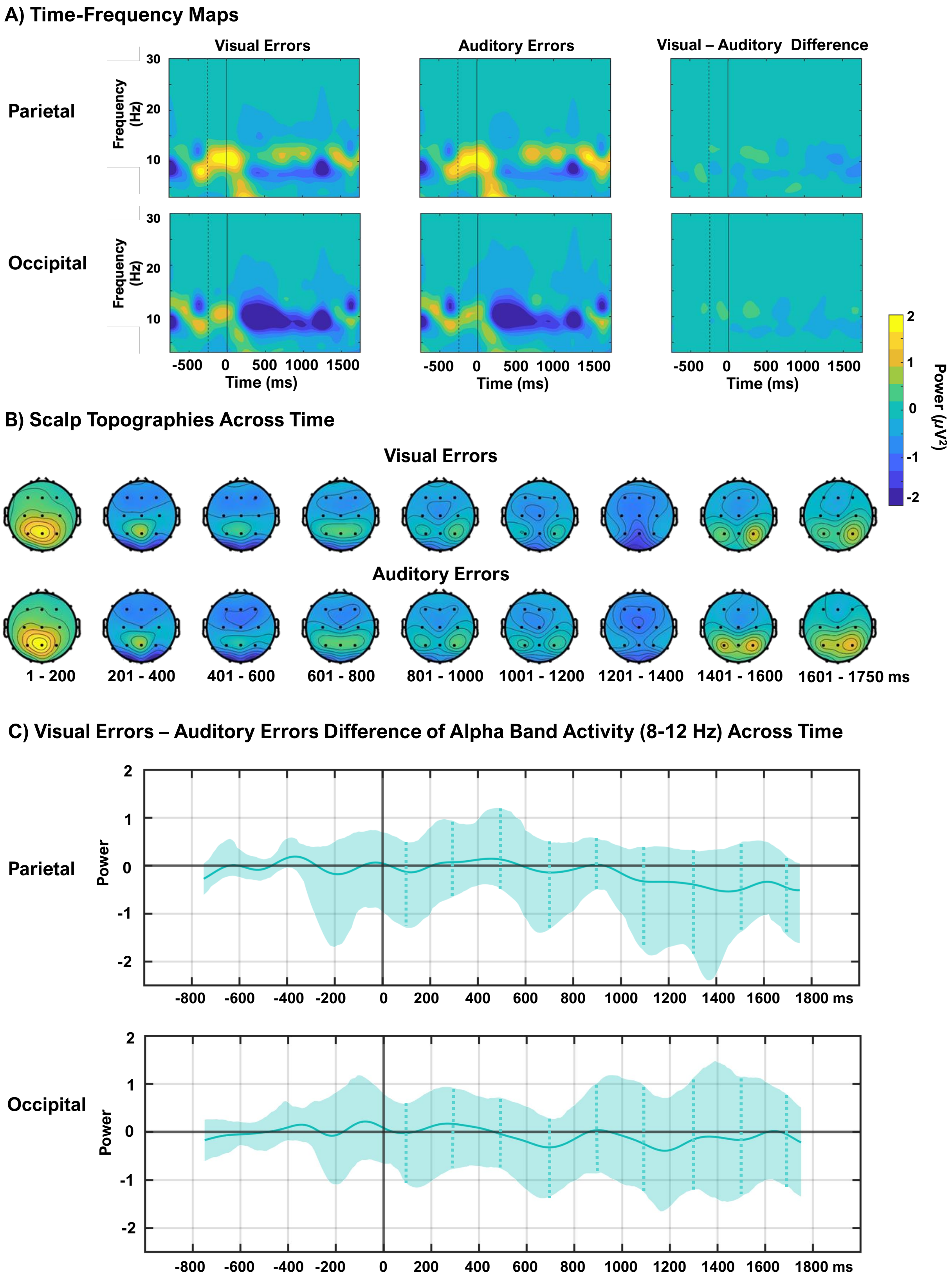

A) Time-Frequency Maps
Visual Errors
Auditory Errors
Visual – Auditory Difference
Parietal
Occipital
Frequency (Hz)
30
20
10
-500
0
500
1000
1500
Time (ms)
Power (µV²)
2
1
0
-1
-2
B) Scalp Topographies Across Time
Visual Errors
Auditory Errors
1 - 200
201 - 400
401 - 600
601 - 800
801 - 1000
1001 - 1200
1201 - 1400
1401 - 1600
1601 - 1750 ms
C) Visual Errors – Auditory Errors Difference of Alpha Band Activity (8-12 Hz) Across Time
Parietal
Occipital
Power
-800
-600
-400
-200
0
200
400
600
800
1000
1200
1400
1600
1800 ms

**Supp. Figure 5**: **Comparison of attend-visual and attend-auditory errors**. Time-frequency maps (A) of the preparatory period for visual error responses, auditory error responses, and the visual minus auditory error response differences. The dotted vertical line indicates the end of the baseline period, the solid vertical line indicates precue onset. Note: statistical testing of the difference map has not occurred, and this panel is displayed for visualization only. Scalp topographies (B) across the preparatory period for visual error trials (top) and auditory error trials (bottom). A and B are on the same color scale. Difference waveforms (C) of the alpha timeseries (8-12 Hz) with 99% bootstrapped confidence intervals indicate no differences in preparation between errors made when attending to either modality. The dotted vertical lines indicate the center of each 200 ms analytic interval. In A and C, the top row includes activity from the parietal electrodes, the bottom includes activity from the occipital electrodes.

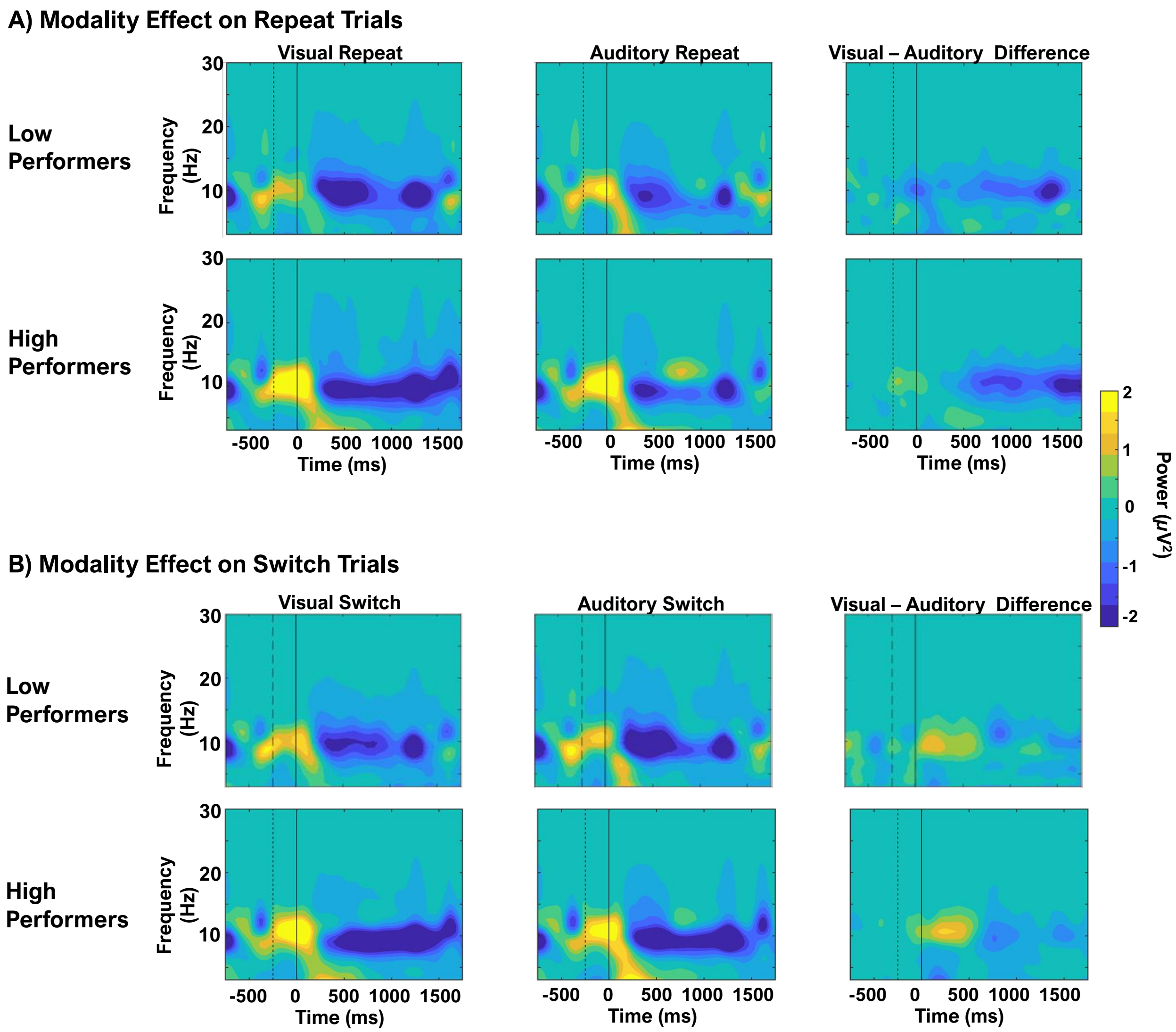


**Supp. Figure 6**: **Modality effects for high and low performers**. Time-frequency maps of the preparatory period for attend-visual repeat responses, attend-auditory repeat responses, and the visual minus auditory response differences (A) for high-performers (top) and low-performers (bottom). Time-frequency maps of the preparatory period for attend-visual switch responses, attend-auditory switch responses, and the visual minus auditory response differences (B) for high-performers (top) and low-performers (bottom). The dotted vertical line indicates the end of the baseline period, the solid vertical line indicates precue onset.

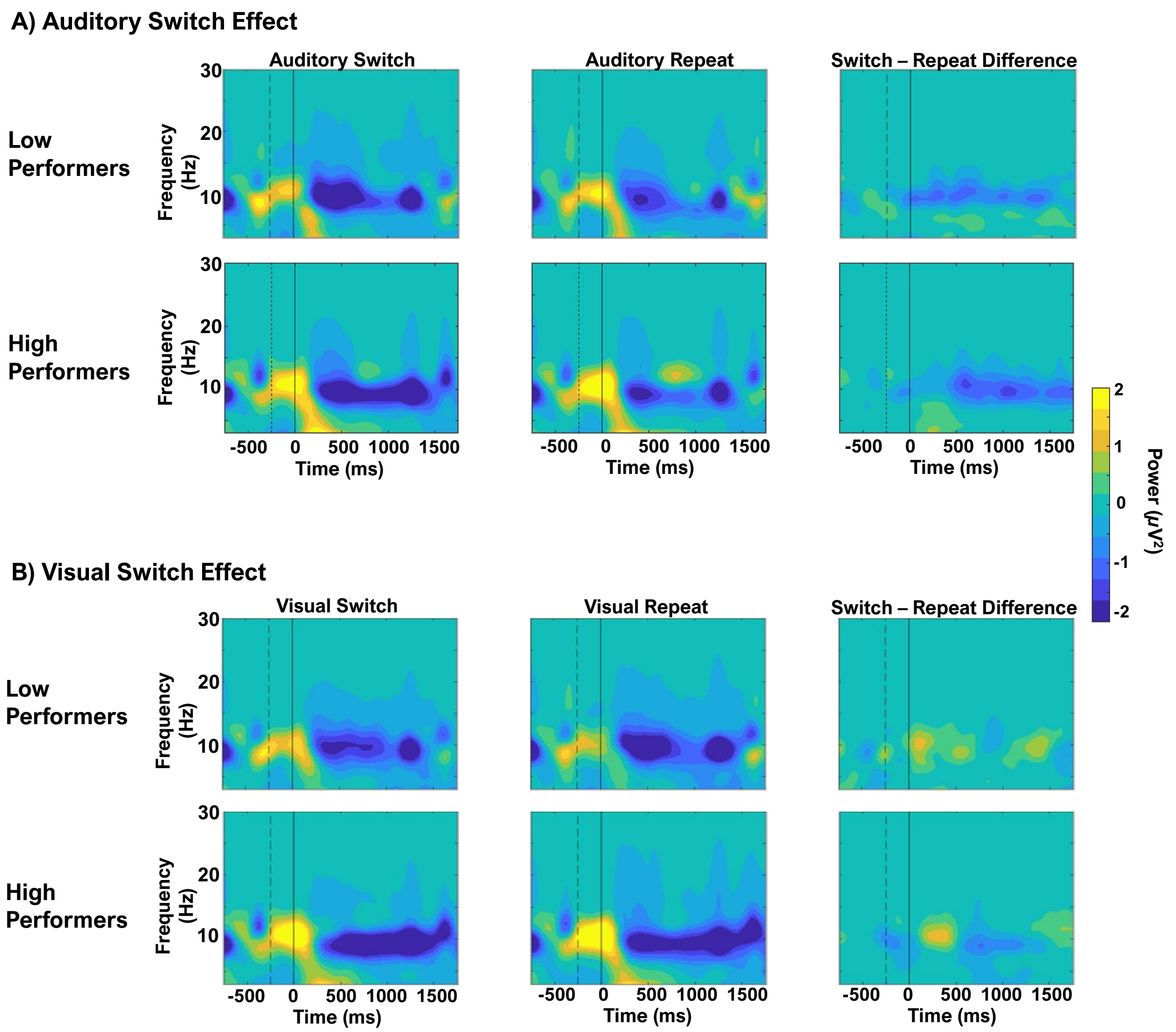


**Supp. Figure 7**: **Switch effects for high and low performers** Time-frequency maps of the preparatory period for attend-auditory switch responses, attend-auditory repeat responses, and the switch minus repeat response differences (A) for high-performers (top) and low-performers (bottom). Time-frequency maps of the preparatory period for attend-visual switch responses, attend-visual repeat responses, and the switch minus repeat response differences (B) for high-performers (top) and low-performers (bottom). The dotted vertical line indicates the end of the baseline period, the solid vertical line indicates precue onset.
